## extended figures for "Phosphoribosyl ubiquitination of SNARE proteins regulate autophagy in Legionella infection"

### Extended Figure EV1

(A)

| Majority protein ID | Gene names | log2 enrichment factor dDup12 over dSidE | -Log10 p-value dDup12 over dSidE | log2 enrichment factor WT over dSidE | -Log10 p-value WT over dSidE |
|---|---|---|---|---|---|
| O00161 | SNAP23 | 4,29 | 2,41 | 3,94 | 2,21 |
| O60749 | SNX2 | 3,72 | 2,87 | 3,97 | 3,17 |
| P56962 | STX17 | 2,39 | 3,17 | 2,47 | 3,45 |
| Q9UNK0 | STX8 | 2,07 | 2,28 | 2,18 | 2,28 |
| O95721 | SNAP29 | 2,96 | 3,27 | 0,94 | 1,89 |
| Q9UNH7 | SNX6 | 1,83 | 2,08 | 2,17 | 2,08 |
| Q13190 | STX5 | 1,60 | 1,64 | 1,72 | 1,83 |

(B)

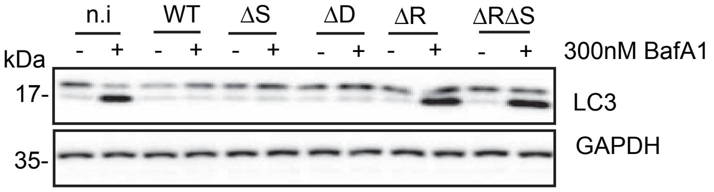


n.i-not infected, WT-wildtype, ΔS-ΔSidE, ΔD-ΔDup, ΔR-ΔRavZ, ΔRΔS-ΔRavZΔSidE

(C)

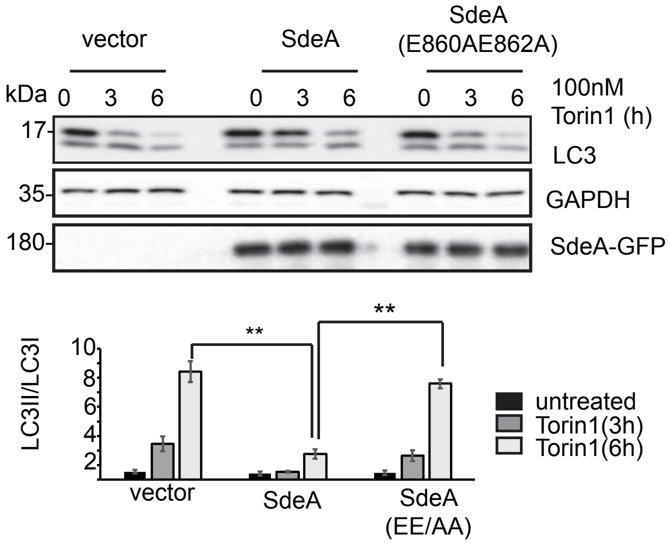


(D)

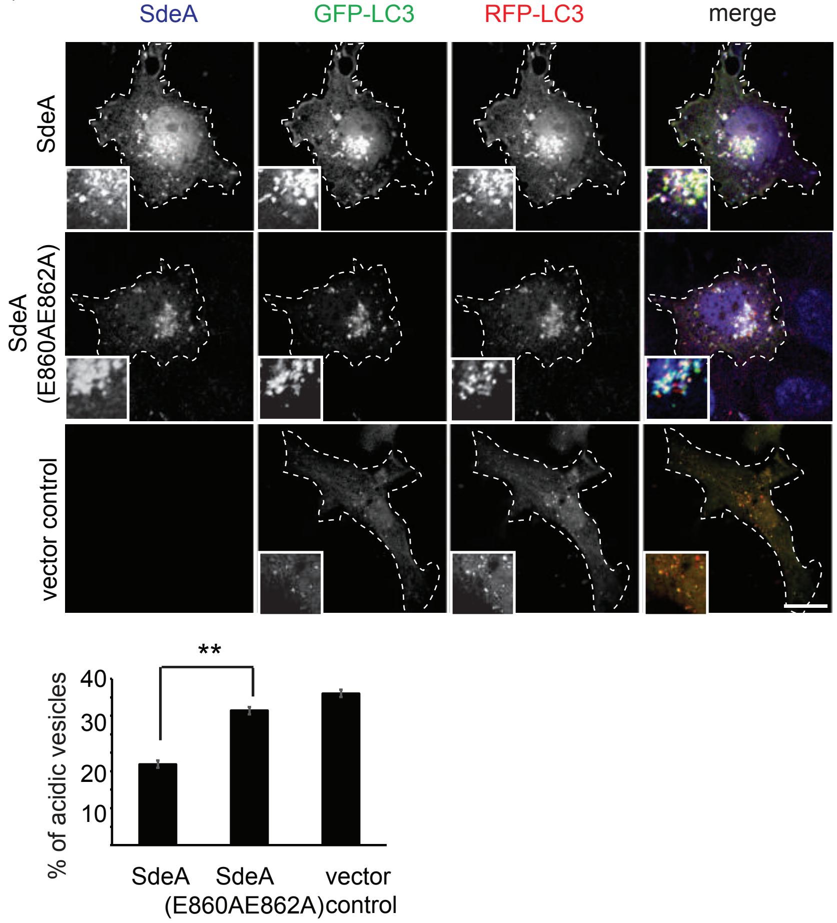


(E)

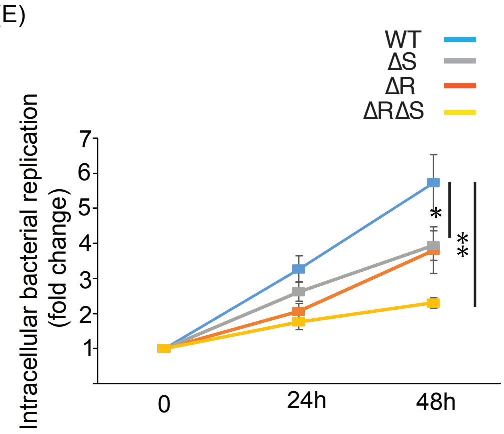


(F)

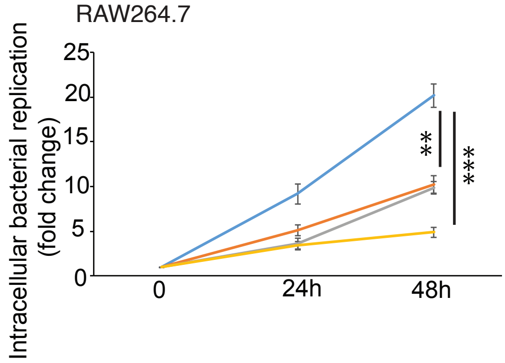

### Extended Figure EV2

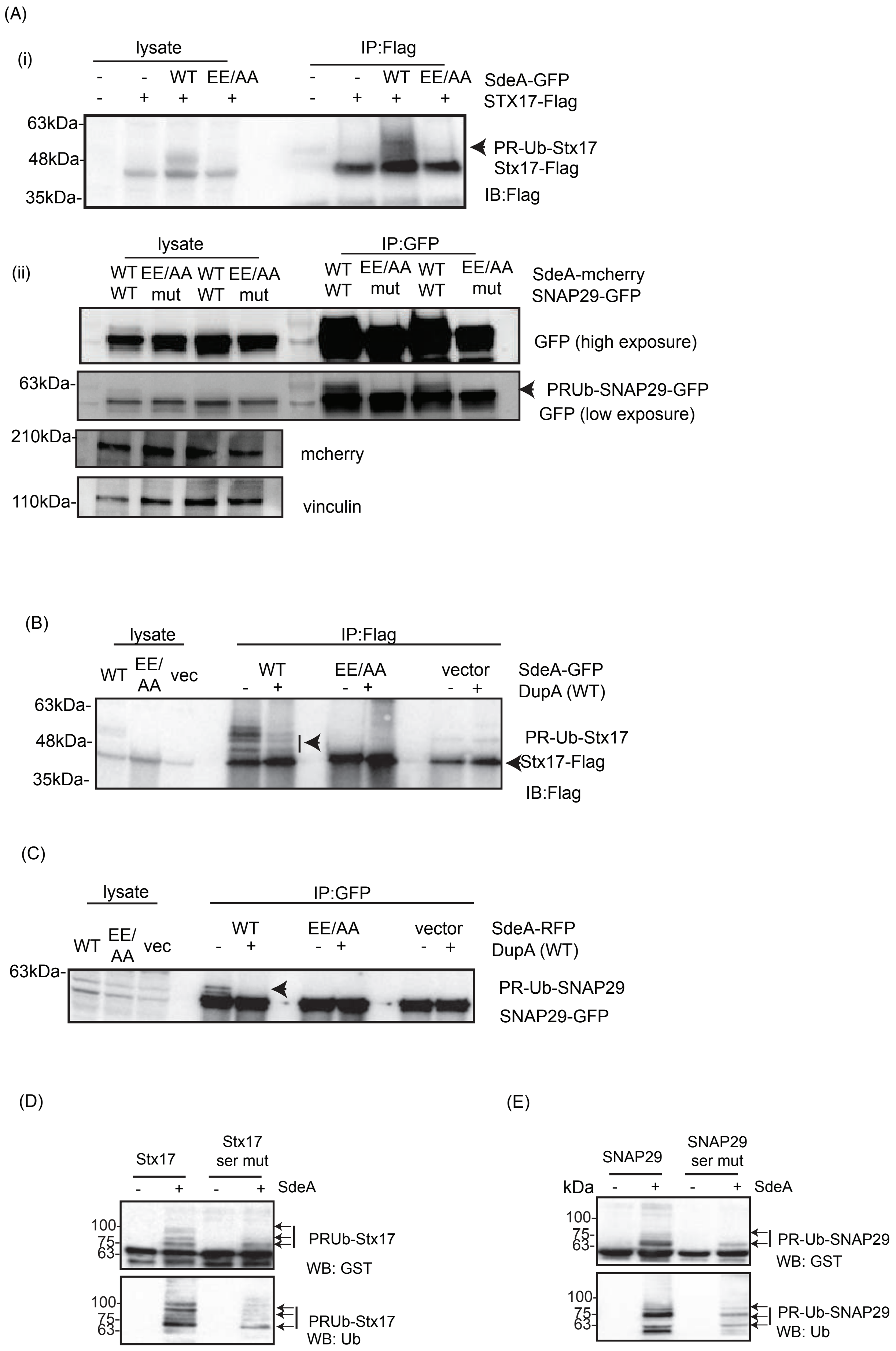

#### Extended Figure EV3

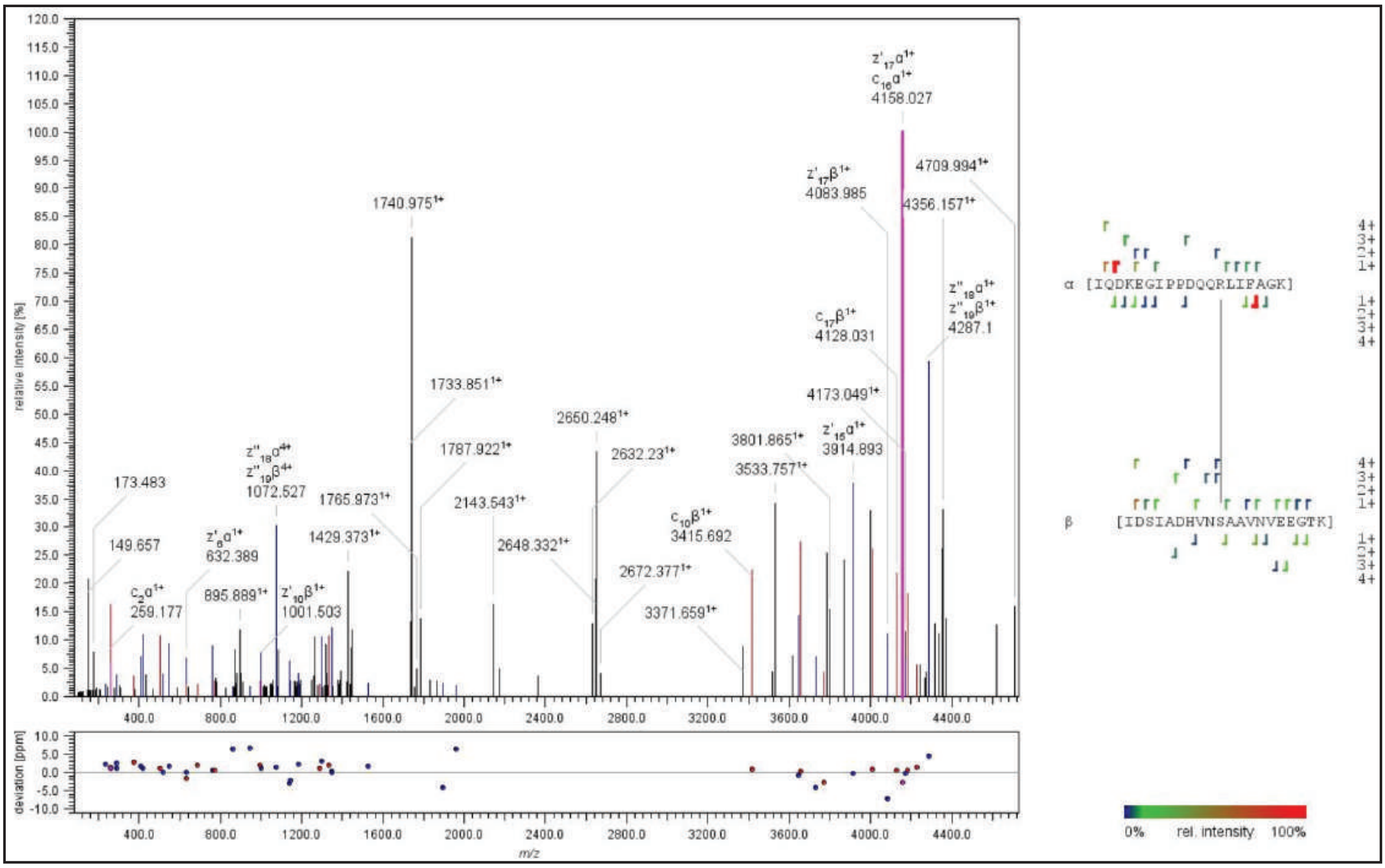


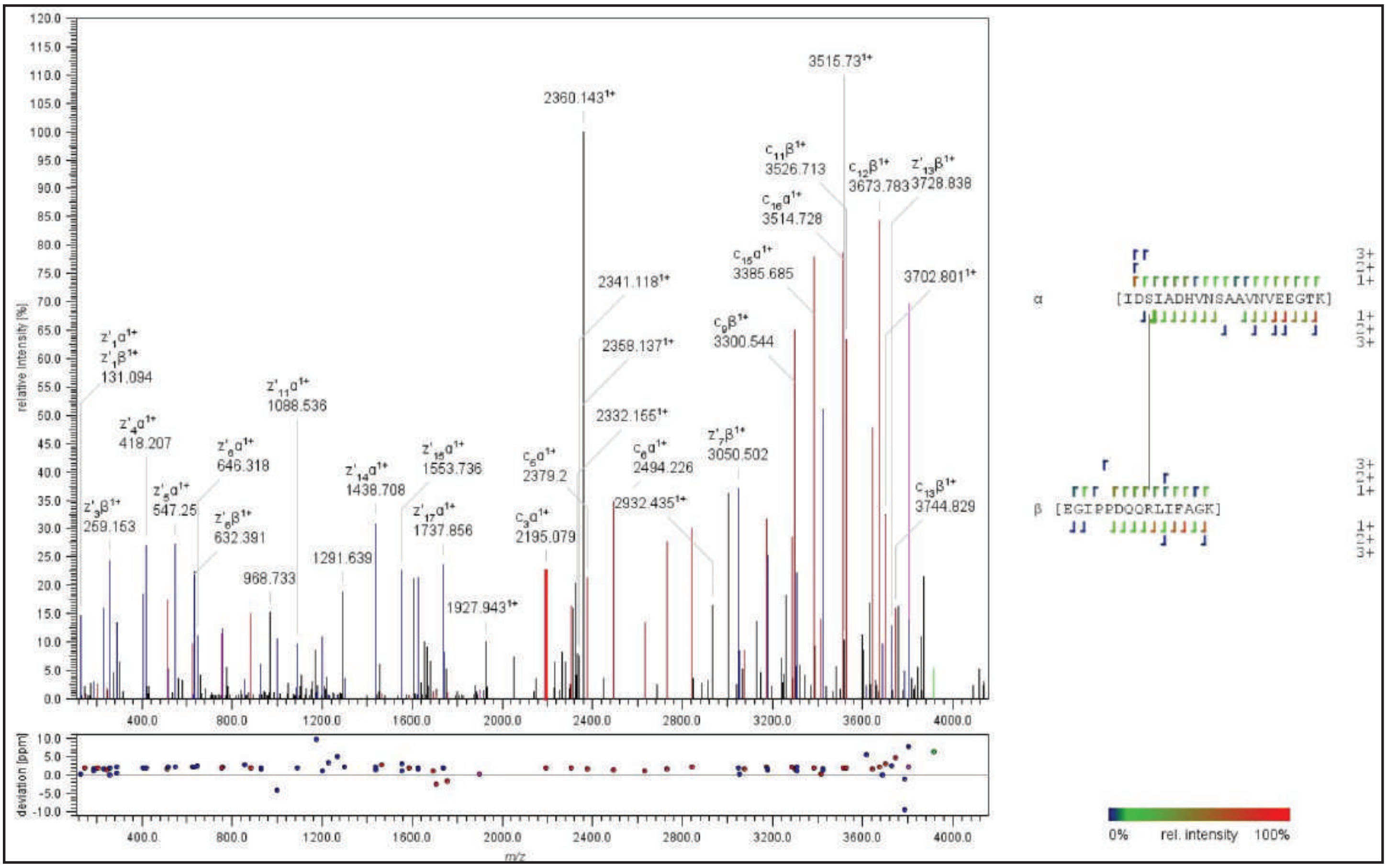


MS spectra of PR-Ub modified STX17(1-224)

### Extended Figure EV4

(A)

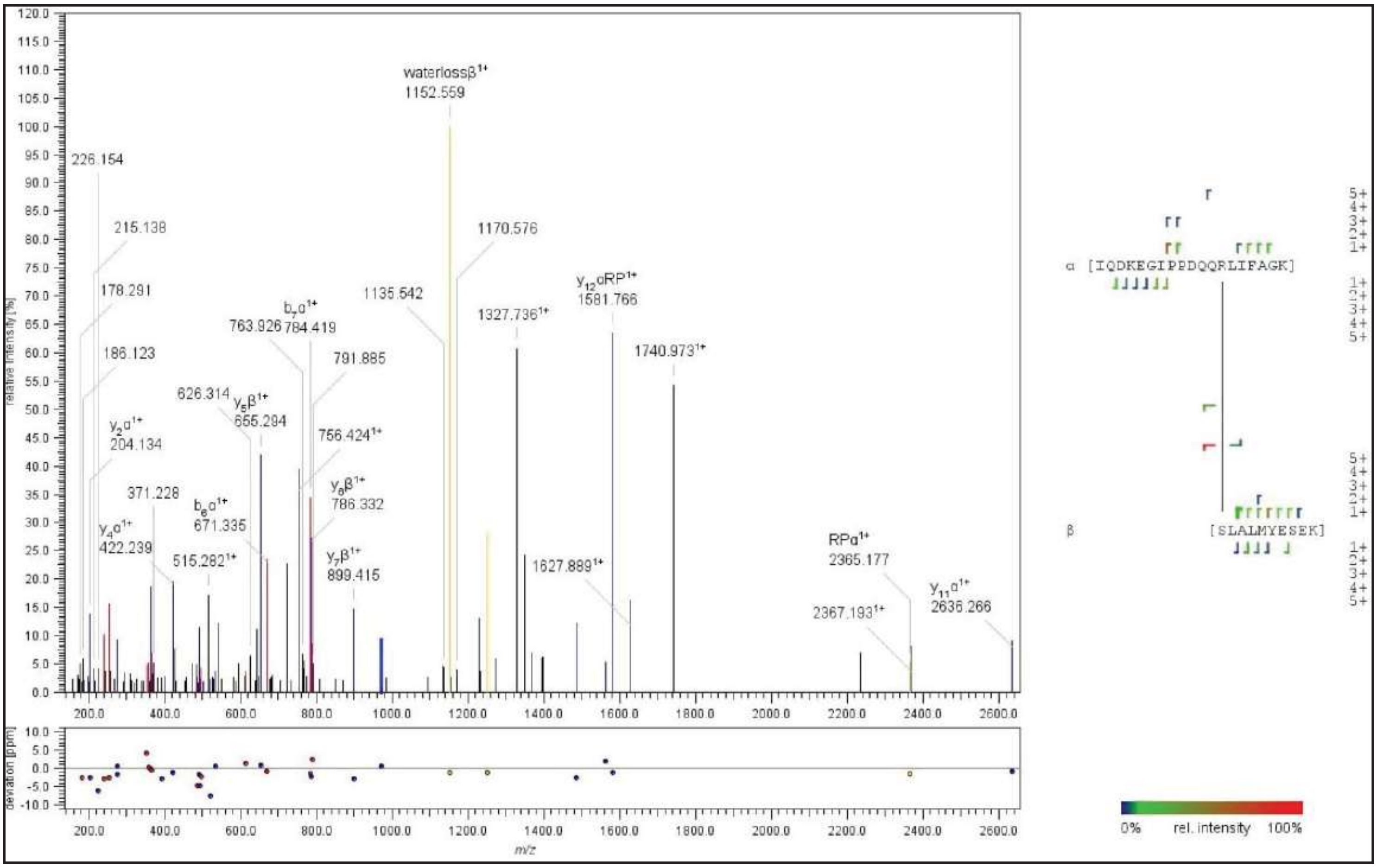


MS spectra of PR-Ub modified SNAP29

(B)

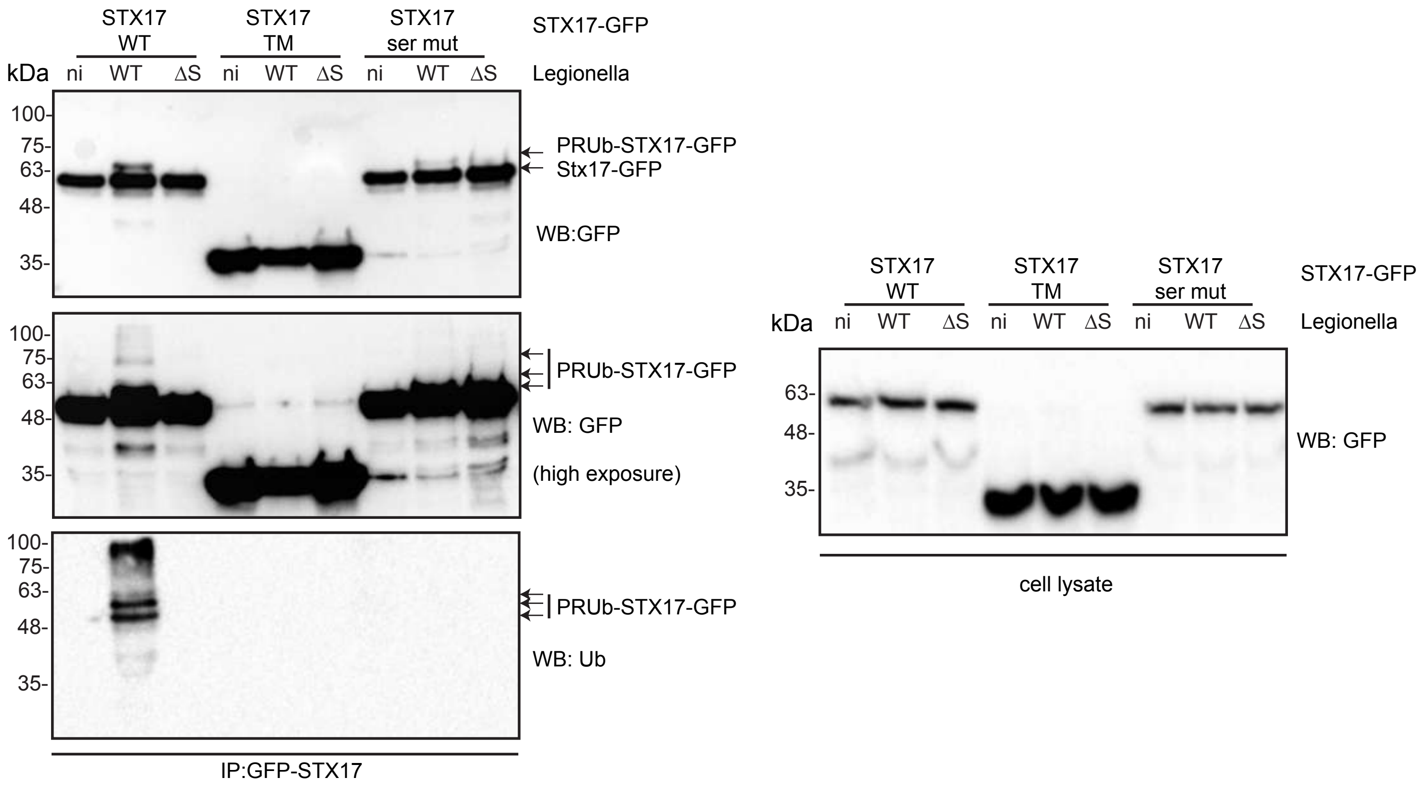

### Extended Figure EV5

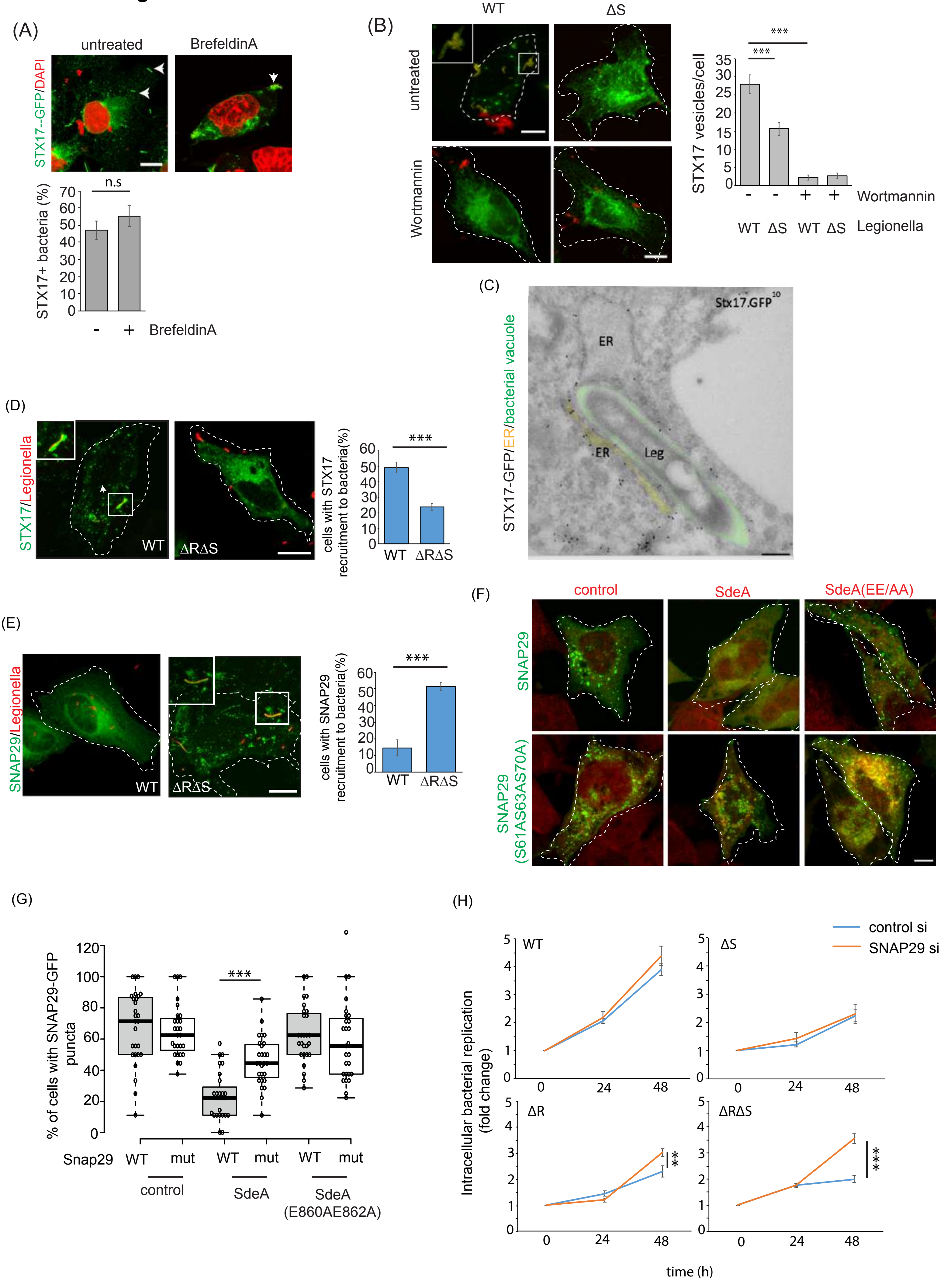

### Extended Figure EV6

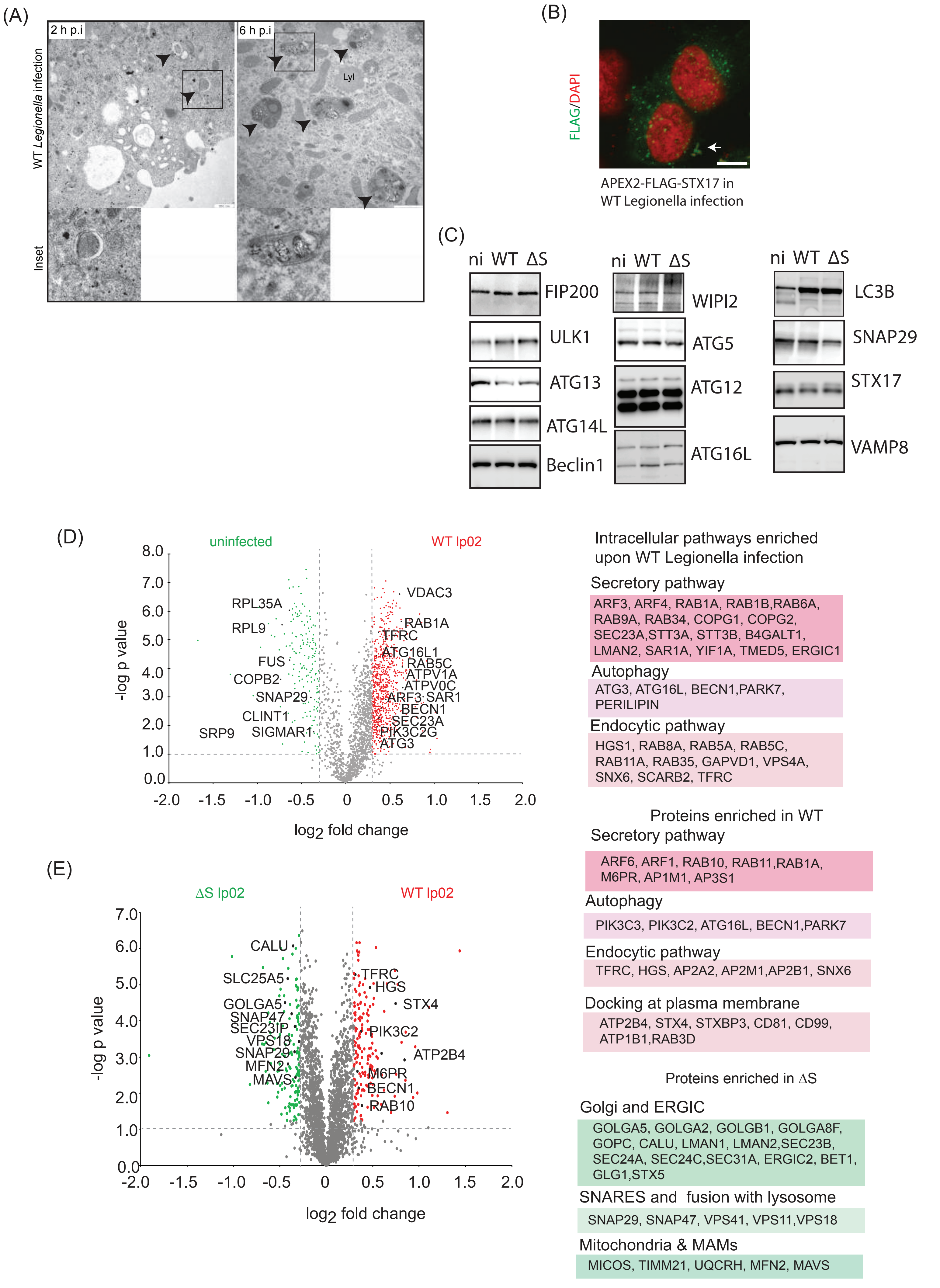
